# High-throughput whole-brain scattering imaging resolves amyloid plaques through clearing-assisted contrast modulation

**DOI:** 10.64898/2026.06.28.735093

**Authors:** Chong Chen, Peilin Gu, Jian Ren

## Abstract

Label-free scattering imaging is widely used in pathology because it enables sensitive tissue assessment without exogenous contrast agents. Yet its limited optical penetration has prevented scattering-based methods from being applied to whole-organ pathology mapping. Here we present clearing-assisted scattering tomography (CAST), a high-throughput, label-free whole-brain mesoscope enabled by selective lipid clearance for scattering enhancement (SELiC). SELiC modulates endogenous refractive-index heterogeneity in cleared tissue, providing whole-brain optical penetration while retaining strong scattering contrast from amyloid plaques and white-matter fibre bundles. CAST enables volumetric imaging of intact mouse brains and brain-wide mapping of amyloid plaque pathology across anatomical regions. This platform establishes a scalable route for label-free, system-level analysis of amyloid pathology and tissue architecture in Alzheimer’s disease (AD) models.

## Introduction

System-scale imaging of intact organs and organisms is emerging as an essential approach in biomedical research, enabling comprehensive spatial mapping of biological structures and disease-associated alterations across whole brains, whole organs and whole bodies^1,2^.Whole-organ imaging is especially important for studies of fiber and vascular connectomics, as these networks span entire biological systems and their connectivity cannot be fully inferred from local fields of view. It is equally important for pathology mapping, where localized measurements are insufficient to capture the regional heterogeneity, spatial gradients and long-range distribution of disease hallmarks^3^.

Nevertheless, systematically charting these vast whole-brain maps remains one of the greatest challenges in today’s biologicial techniques, due to their unprecedented scale and complexity^4^. This effort requires a comprehensive 3D visualization that can see the big (i.e., the entire brain) and the small (i.e., individual Aβ plaques). Fundamentally, this imposes a demand for imaging systems to have high spatial throughput. In addition, the imaging procedure also needs to have a high temporal throughput, shortening the overall turnaround time of the above whole-brain mapping by a low-cost and less labor-intensive workflow, thus making it more adoptable for large-cohort longitudinal studies covering multiple stages of Alzheimer’s disease (AD) progression.

Towards these demands, numerous imaging efforts have been made spanning multiple scales with various modalities. Positron emission tomography (PET), the current clinical gold standard for detecting Aβ pathology, enables in vivo whole-brain visualization and quantitative mapping of Aβ deposition using Aβ-binding radiotracers^5^. However, its limited spatial resolution precludes detailed characterization of plaque-associated microstructural pathology. Currently, light microscopy, such as fluorescent micro-optical sectioning tomography (fMOST)^6^, enables whole-brain mapping of Aβ distribution with a 1∼2 μm resolution by tissue sectioning to overcome the light penetration limit. However, tissue sectioning disrupts samples’ structural integrity and significantly reduces imaging temporal throughput, limiting LM’s further applications to longitudinal AD studies.

Recently, benefiting from tissue clearing’s advancement, it is possible to image through thick tissues and even intact murine brains^7–10^. Meanwhile, light-sheet microscopy (LSM) allowing for convenient volumetric imaging of transparent specimens^11,3,12,4^. Yet, the limited diffusion rate of macromolecules (i.e., antibodies) significantly prolongs the fluorescence labeling process for intact brain tissue. The characteristic diffusion time *t* for a molecule to penetrate a tissue over a distance *L* is governed by the equation *t≈L*^2^/2D, where *D* is the diffusion coefficient, indicating that the labeling time increases quadratically with tissue thickness^13^. Complete penetration of IgG antibodies into an intact adult mouse brain would require approximately 3-4 weeks^14,15^. While this duration might be feasible for some small-scale studies, the quadratic dependency will make labeling large tissue samples very difficult. For example, achieving uniform antibody diffusion in an intact macaque brain would require approximately 6 to 8 months, and a full antibody-labeling of a human brain specimen could take over 5 years, posing an astonishing barrier that makes human whole-brain Aβ immunofluorescence imaging virtually unattainable at present. In addition, the finite fluorescence yield limits imaging throughput and places a practical constraint on large samples^16,17^. While numerous instrument advancements have been made for LSM, regardless of the hardware techniques employed, the throughput of fluorescence imaging is fundamentally limited by the minimal dwell time determined by the amount of photons emitted from fluorophores. Even with the brightest commercial fluorescence dye (i.e., Alexa dye), it still takes a great number of hours or days to scan a whole adult mouse brain. The acquisition time quickly becomes impractical when extrapolated to larger models. For example, it would require an uninterrupted period of 12 days for state-of-the-art LSMs to image a macaque brain at a resolution of 1 μm^3^.^18^ For a human brain, this number grows to a staggering 3 months or more.

To address these challenges, we present a high-throughput, label-free whole-brain mesoscope,named CAST, enabled by selective lipid clearance for scattering enhancement (SELiC). CAST captures elastic scattering from intrinsic Aβ pathology, providing long-lasting and photostable contrast for AD-related biomarkers. From a fundamental physics perspective, this elastic scattering process conserves the photons’ and the system’s energies, ensuring its time invariance (infinitely repeatable) and exempting it from photobleaching, a consequence of energy dissipation. Moreover, CAST imaging exploits tissue intrinsic contrast; therefore, no labeling is required. Bypassing the diffusion barrier, this new imaging approach would greatly decrease tissue preparation time and cost, allowing for an easy-to-adopt and less labor-intensive workflow for large-cohort AD studies.

## Results

### CAST system and validation

Whole-organ pathology mapping requires an imaging system that combines a long working distance, cellular-scale resolution and a centimetre-scale field of view. This combination is difficult to achieve with conventional off-the-shelf microscope objectives, which typically trade FOV, numerical aperture and working distance against one another^19,20^. Recent high-throughput optical imaging systems have addressed this challenge using custom-designed objectives, but such solutions are often costly and not easily adaptable across platforms^21–23^. To overcome this limitation, we adapted consumer photographic lenses for CAST imaging. Owing to their large image circle, long flange focal distance and high light-collection capacity, photographic lenses provide a practical route to large-field coherent detection. Similar design principles have recently been used in mesoscopic light-sheet microscopy, including mesoSPIM, to image cleared centimetre-scale specimens with micrometre-scale resolution^19^.

CAST imposes more demanding optical requirements than light-sheet microscopy because it operates in the near-infrared/SWIR regime and requires broadband detection (**Fig.1a**)^24^. After extensive optical evaluation and experimental testing, we selected a SWIR-optimized photographic lens (SWIR-16, Navitar Inc.) for the CAST detection path. With an F/1.4 aperture, a 13.5-mm working distance in air and a field of view exceeding 15 mm, this lens provided the combination of light-collection efficiency, working distance and centimetre-scale coverage required for whole-organ CAST imaging. To fully exploit the high etendue of the SWIR-16 lens, we incorporated a paired off-axis parabolic mirror (MPD239-P0, Thorlabs Inc.) relay into the scanning path. The relay positioned the scanning galvanometer at a plane conjugate to the lens pivot point, preserving optical throughput and maintaining stable large-field scanning across the centimetre-scale FOV. As shown in Fig. 1a, the detection system of CAST was implemented using optical frequency-domain imaging, also known as coherent frequency-domain reflectometry. The system was driven by a custom-built wavelength-tunable external-cavity laser (WT-ECL), which was swept at 50 kHz over a 20-THz optical bandwidth centred at 1.3 μm, with an instantaneous linewidth of 34 GHz^25^.

**Fig 1.**
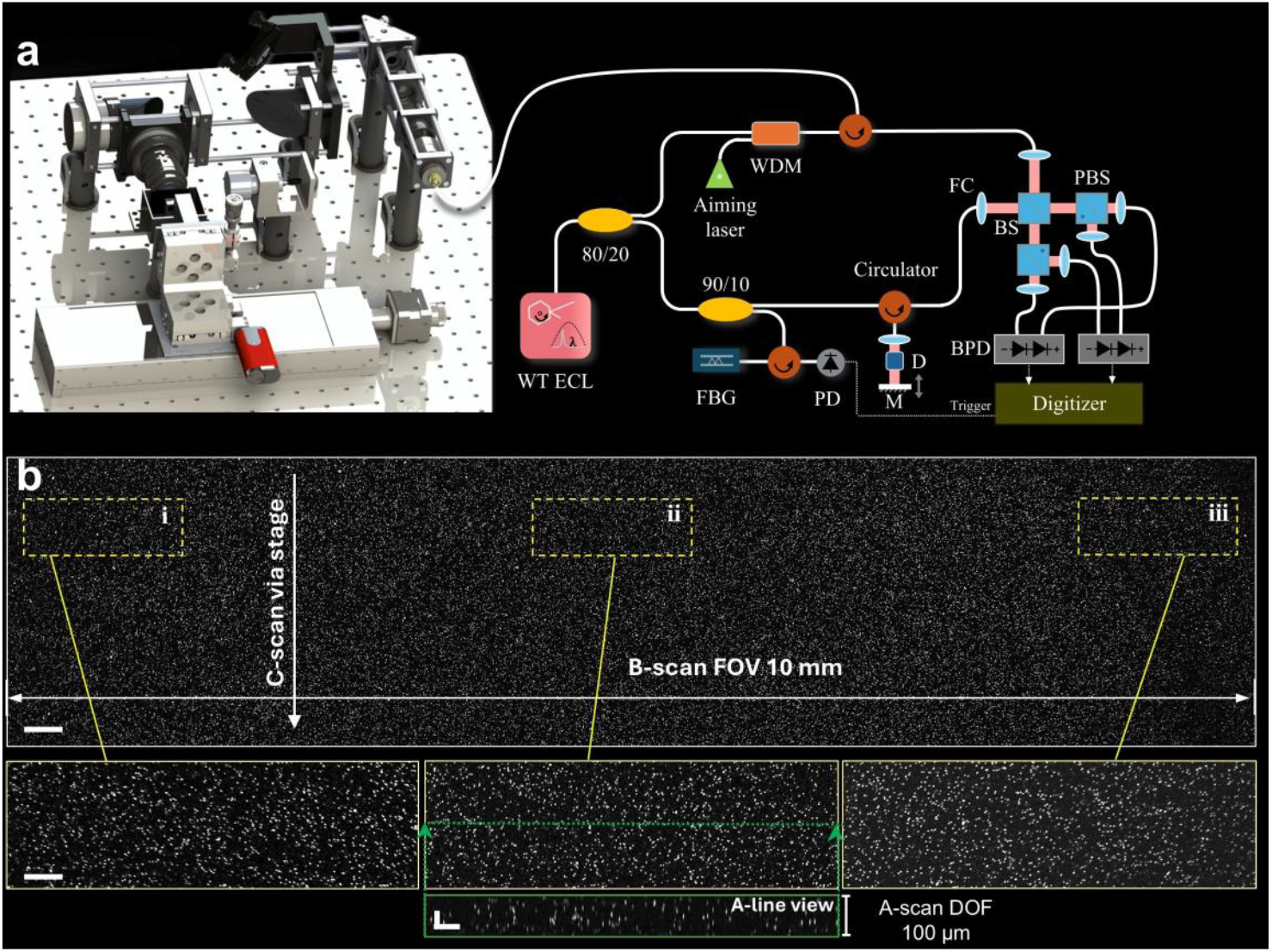
CAST platform for rapid micrometre-scale imaging of intact whole brains. **a**, Schematic of the CAST system, integrating a high-throughput large-field scanning module based on a photographic lens with a high-sensitivity coherent detection unit. **b**, Microsphere phantom characterization of the CAST imaging performance. The system achieved an approximately 3-μm lateral resolution over a 10-mm scanning field of view at a central wavelength of 1.3 μm, sufficient to cover the dorsoventral extent of an adult mouse brain, with an axial resolution of approximately 6 μm in water.

To characterize the optical performance of the CAST system, we prepared a microsphere phantom by embedding 500-nm beads (115073, Lab261 Inc.) in an 8% acrylamide hydrogel. CAST imaging of this phantom showed that the system achieved an approximately 3-μm lateral resolution across a 10-mm field of view and an approximately 6-μm axial resolution. The field of view along the slow-scan direction was primarily determined by the travel range of the translation stage, allowing scalable extension of the imaging volume. This optical field of view is sufficient to span the dorsoventral extent of an adult mouse brain in a single scan.

### Multiplexed CAST imaging of Aβ plaques and fiber bundles using SELiC clearing method

Native tissues strongly scatter light, limiting the penetration depth of scattering-based optical imaging and increasing the contribution of multiple scattering artifacts^26^. Tissue clearing mitigates this problem by homogenizing refractive-index variations across tissue constituents^27^. However, this process is not uniform: although hydration and refractive-index matching can reduce scattering from protein-rich structures, hydrophobic lipids can hinder water penetration and prevent complete refractive-index equilibration (Fig.2a). This creates a competing requirement for CAST imaging, in which lipid-associated background must be suppressed while diagnostically useful endogenous scattering from protein-rich pathological and fibre structures must be retained.

To address this challenge, we developed selective lipid clearance for scattering enhancement (SELiC) (Fig.2b). Fixed brains were further treated with 1% glutaraldehyde (GA) to enhance tissue stabilization. GA-mediated crosslinking preserves protein-rich tissue architecture and decreases tissue permeability, which can limit lipid extraction in lipid-rich regions such as white matter. This effect creates a useful contrast mechanism for CAST: grey-matter regions can be sufficiently cleared to reduce background scattering, whereas lipid-rich fibre bundles retain residual refractive-index heterogeneity and remain strongly visible. Following SELiC, samples were embedded in 16% acrylamide gel and immersed in refractive-index-matching buffer to facilitate stable mounting and CAST imaging.

In Fig. 2c, plaque-like structures in APP/PS1 mouse brains were clearly visualized across multiple brain regions, whereas similar structures could not be found in the control animal (Fig. 2d). The magnified view of the cortical region reveals that these plaque structures exhibit significantly stronger scattering than the surrounding tissue. To validate the plaque-like structures detected by CAST, these structures were subsequently co-localized with the findings of the same brain slice, but from fluorescence microscopy with 6E10 stainning. The highly similar spatial patterns confirmed that the structures visualized by CAST imaging correspond to Aβ. CAST provides an efficient strategy for brain-wide mapping of Aβ pathology. Unlike conventional fluorescence-based approaches that require prolonged staining and labelling procedures, CAST enables whole-brain imaging after tissue clearing alone, allowing label-free detection of plaque-associated scattering signals across intact brains. Beyond Aβ pathology, CAST also resolved mesoscale brain-wide fiber bundle architecture (Fig. 2d). Magnified views further demonstrated clear visualization of fiber structures across distinct anatomical regions, highlighting the ability of CAST to capture both pathological scattering signals and endogenous structural contrast within the same intact brain.

**Fig 2.**
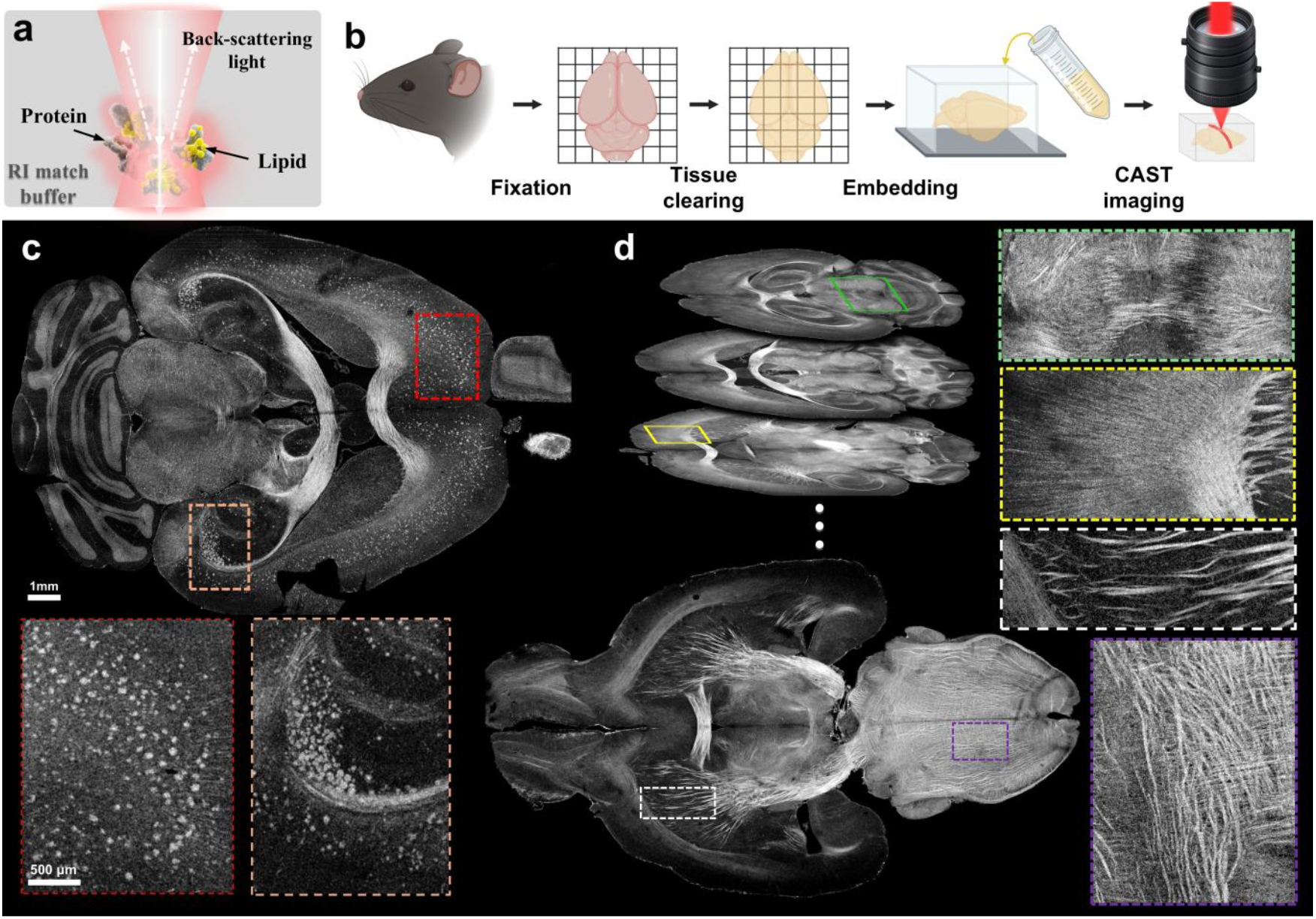
CAST workflow for whole-brain volumetric imaging. **a**, Schematic illustrating the origin of CAST contrast in cleared tissue, where residual refractive-index heterogeneity among proteins, lipids and the matching buffer generates optical scattering. **b**, Workflow of selective lipid clearance for scattering enhancement (SELiC), which reduces lipid-associated background scattering while preserving protein-rich endogenous contrast for CAST imaging. **c**, CAST imaging of a SELiC-treated APP-PS1 Alzheimer’s disease mouse brain showing prominent scattering signals associated with Aβ plaques. d, SELiC enables controlled lipid clearance in white-matter regions, enabling clear identification of brain-wide fiber bundle scattering signals.

### Aβ plaque segmentation and brain-wide pathology mapping

To quantify Aβ pathology across the whole brain, we established an integrated analysis pipeline that combines deep-learning-based plaque segmentation with CAST-tailored registration to a reference brain atlas. This workflow enabled automated detection of plaque-associated scattering signals and their anatomical assignment to defined brain regions (Fig.3).

Because CAST contrast is generated by residual refractive-index heterogeneity in lipid-retaining structures, Aβ plaque-associated signals coexist with endogenous fibre-bundle contrast in the same image volume (Fig. 3a). Intensity alone was therefore insufficient for robust plaque identification. We built an optimized U-Net segmentation network (Fig.3b) using ground-truth labels generated by three independent researchers from 100 manually annotated axial brain images. Overlay analysis showed that the segmented plaque masks closely matched plaque-associated bright scattering signals in the raw CAST data (Fig. 3d), supporting the reliability of the network output. The resulting brain-wide 3D rendering revealed regionally heterogeneous Aβ plaque densities (Fig. 3c), in agreement with previous whole-brain imaging studies of Aβ deposition patterns in APP/PS1 mice.

**Fig 3.**
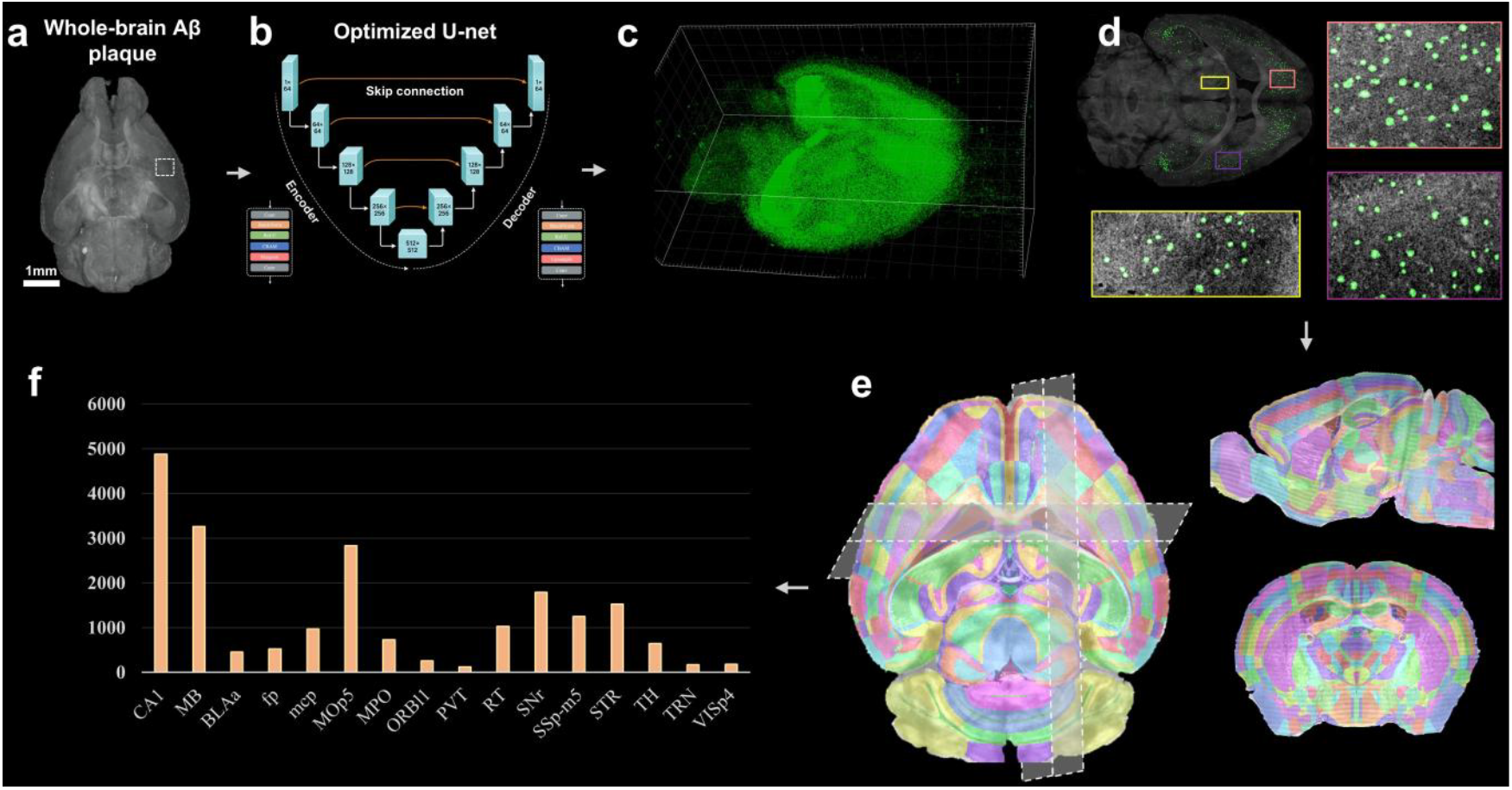
Brain-wide mapping of Aβ pathology in an Alzheimer’s disease mouse model. **a**, Maximum-intensity projection rendering of brain-wide Aβ plaque-associated scattering signals in an APP-PS1 mouse brain. The white box indicates a representative training volume used for deep-learning-based plaque segmentation. **b**, Whole-brain CAST data are passed to optimized U-net segmentation module to generate binary segmentation maps. **c**, Three-dimensional rendering of deep-learning-segmented Aβ plaque distribution across the whole brain. **d**, Overlay of raw CAST data and segmented Aβ plaque masks in a representative mid-brain axial section. Magnified views show that the green segmentation masks closely overlap with plaque-associated bright scattering signals. **e**, Anatomical registration of CAST datasets to the Allen Reference Atlas (ARA) using a custom elastix-based pipeline, enabling brain-wide localization of segmented Aβ plaques. **f**, Atlas-based quantification of Aβ plaque number across brain regions.

As no reference atlas exists for scattering-based brain images, we established a CAST-specific atlas-registration pipeline. CAST fibre-bundle contrast showed strong structural correspondence with an intensity-inverted Allen Reference Atlas (ARA), enabling us to construct an inverted ARA template for registration. Manually placed anatomical landmarks were first used to initialize rigid and affine transformations, followed by elastix-based non-linear registration. Orthogonal overlays of the registered atlas annotations and raw CAST data showed good anatomical agreement across brain regions (Fig. 3e), enabling brain-wide assignment of segmented Aβ plaques to atlas-defined regions. Regional quantification across 16 representative brain regions revealed heterogeneous Aβ plaque burdens throughout the brain (Fig. 3f).

## Acknowledgements

This research was funded by National Institutes of Health (NIH) (R00AG059946)

## Discussion

Here we present CAST, a high-throughput, label-free scattering tomography platform for simultaneous whole-brain imaging of Aβ plaques and white-matter fibre bundles. By repurposing photographic lenses as large-field scanning optics, CAST achieves cellular-scale volumetric imaging across intact mouse brains without fluorescence labelling. In parallel, we developed selective lipid clearance for scattering enhancement (SELiC), which enables whole-brain optical penetration while retaining high endogenous scattering contrast from lipid-rich pathological and fibre structures. This combination establishes CAST as a powerful tool for brain-wide Aβ pathology mapping and mesoscale fibre-architecture analysis. To support quantitative pathology mapping, we further developed a complete analysis workflow that integrates optimized deep-learning-based Aβ plaque segmentation with a CAST-specific atlas-registration method for regional whole-brain quantification.

The throughput of CAST has substantial room for further improvement. Our current system used a custom-built 50-kHz swept source to demonstrate whole-brain label-free scattering tomography, but the acquisition speed of coherent scattering imaging can scale directly with the A-line rate. Swept-source OCT has already reached multi-MHz to 160-MHz sweep rates in research systems, and 10-MHz-class sources have recently been demonstrated for biomedical imaging, indicating that CAST could benefit immediately from faster light sources^28^. In principle, source-speed scaling alone could reduce whole-brain CAST acquisition to the order of tens of seconds at ∼3-μm sampling resolution. Beyond light-source speed, the current Gaussian-beam design uses only a limited axial fraction of each A-line because of its finite depth of focus. Incorporating extended-depth-of-focus illumination, multifocal imaging or computational adaptive optics could increase the usable axial range per scan and further improve CAST throughput by orders of magnitude.

Beyond Aβ plaques, CAST also provides robust endogenous contrast for white-matter fibre bundles. Its high-throughput, whole-brain imaging capability makes it well suited for large-cohort studies aimed at examining AD-stage-dependent alterations in mesoscale fibre architecture and connectome-level organization.

## References

1. Richardson, D. S. & Lichtman, J. W. Clarifying tissue clearing. Cell 162, 246–257 (2015).

2. Mai, H. et al. Scalable tissue labeling and clearing of intact human organs. Nat Protoc 17, 2188–2215 (2022).

3. Glaser, A. K. et al. A hybrid open-top light-sheet microscope for versatile multi-scale imaging of cleared tissues. Nat Methods 19, 613–619 (2022).

4. Glaser, A. et al. Expansion-assisted selective plane illumination microscopy for nanoscale imaging of centimeter-scale tissues. eLife 12, (2025).

5. Chapleau, M., Iaccarino, L., Soleimani-Meigooni, D. & Rabinovici, G. D. The Role of Amyloid PET in Imaging Neurodegenerative Disorders: A Review. J Nucl Med 63, 13S–19S (2022).

6. Zhong, Q. et al. High-definition imaging using line-illumination modulation microscopy. Nat Methods 18, 309–315 (2021).

7. Chung, K. et al. Structural and molecular interrogation of intact biological systems. Nature 497, 332–337 (2013).

8. Zhao, S. et al. Cellular and Molecular Probing of Intact Human Organs. Cell 180, 796–812.e19 (2020).

9. Murakami, T. C. et al. A three-dimensional single-cell-resolution whole-brain atlas using CUBIC-X expansion microscopy and tissue clearing. Nat Neurosci 21, 625–637 (2018).

10. Susaki, E. A. et al. Versatile whole-organ/body staining and imaging based on electrolyte-gel properties of biological tissues. Nat Commun 11, 1982 (2020).

11. Voigt, F. F. et al. The mesoSPIM initiative: open-source light-sheet microscopes for imaging cleared tissue. Nat Methods 16, 1105–1108 (2019).

12. Patel, K. B. et al. High-speed light-sheet microscopy for the in-situ acquisition of volumetric histological images of living tissue. Nat. Biomed. Eng 6, 569–583 (2022).

13. Ku, T. et al. Elasticizing tissues for reversible shape transformation and accelerated molecular labeling. Nat Methods 17, 609–613 (2020).

14. Renier, N. et al. iDISCO: A Simple, Rapid Method to Immunolabel Large Tissue Samples for Volume Imaging. Cell 159, 896–910 (2014).

15. Kirst, C. et al. Mapping the fine-scale organization and plasticity of the brain vasculature. Cell 180, 780–795.e25 (2020).

16. Murray, J. M., Appleton, P. L., Swedlow, J. R. & Waters, J. C. Evaluating performance in three-dimensional fluorescence microscopy. J Microsc 228, 390–405 (2007).

17. Joshi, P., Kumar, P., s, A., Varghese, J. M. & Mondal, P. P. Fluorescence-based multifunctional light sheet imaging flow cytometry for high-throughput optical interrogation of live cells. Commun Phys 7, 25 (2024).

18. Xu, F. et al. High-throughput mapping of a whole rhesus monkey brain at micrometer resolution. Nat Biotechnol 39, 1521–1528 (2021).

19. Vladimirov, N. et al. Benchtop mesoSPIM: a next-generation open-source light-sheet microscope for cleared samples. Nat Commun 15, 2679 (2024).

20. Glaser, A. et al. Expansion-assisted selective plane illumination microscopy for nanoscale imaging of centimeter-scale tissues. eLife 12, (2024).

21. Fan, J. et al. Video-rate imaging of biological dynamics at centimetre scale and micrometre resolution. Nat. Photonics 13, 809–816 (2019).

22. Yang, M. et al. Ultra-wide-field, deep, adaptive two-photon microscopy for multi-scale neuronal imaging. Light Sci Appl 15, 198 (2026).

23. Yu, C.-H. et al. The Cousa objective: a long-working distance air objective for multiphoton imaging in vivo. Nat Methods 21, 132–141 (2024).

24. Ren, J., Choi, H., Chung, K. & Bouma, B. E. Label-free volumetric optical imaging of intact murine brains. Sci Rep 7, 46306 (2017).

25. Lippok, N. et al. Depolarization signatures map gold nanorods within biological tissue. Nature Photon 11, 583–588 (2017).

26. Untracht, G. R. et al. Spatially offset optical coherence tomography: Leveraging multiple scattering for high-contrast imaging at depth in turbid media. Science Advances 9, eadh5435 (2023).

27. Richardson, D. S. et al. Tissue clearing. Nat Rev Methods Primers 1, 1–24 (2021).

28. Zhu, S. et al. 160 MHz optical coherence tomography for highly scattering biomedical imaging. Photon. Res., PRJ 14, 2980–2996 (2026).

